# Sphingolipid regulation by yeast Mdm1 supports adaptive remodeling of the methionine transporter Mup1

**DOI:** 10.64898/2026.02.25.707973

**Authors:** Daniel Adebayo, Eseiwi Obaseki, Kashvi Vasudeva, Marwa Aboumourad, Scott Miller, Anne Ostermeyer-Fay, Daniel Canals, Xun Bao, Jing Li, Hanaa Hariri

## Abstract

Membrane lipid composition influences endocytic remodeling of nutrient transporters, yet how lipid metabolism is spatially coordinated to support sustained adaptation to nutrient limitations remains unclear. Here, we investigated whether the ER-vacuole tether Mdm1 links sphingolipid homeostasis to regulation of the high-affinity methionine permease Mup1 in budding yeast. To test this, we examined Mup1 trafficking, amino acid homeostasis, and sphingolipid composition in *mdm1*Δ cells during starvation. We found that loss of Mdm1 causes persistent retention of Mup1 at the plasma membrane, accompanied by reduced intracellular methionine and broad amino acid depletion. Lipidomic analyses revealed decreased sphingoid bases and altered ceramide composition in *mdm1*Δ cells. Importantly, supplementation with the sphingolipid precursor phytosphingosine restored sphingolipid pools, rescued Mup1 endocytosis, and improved amino acid homeostasis. Consistent with a chronic amino acid restriction-like state, *mdm1*Δ cells exhibited extended chronological lifespan. Together, these findings identify Mdm1 as a spatial organizer of sphingolipid metabolism required for adaptive endocytic remodeling of Mup1, thereby linking ER-vacuole contact site function to plasma membrane proteostasis and metabolic adaptation.

## Introduction

Eukaryotic cells adapt to changes in nutrient fluctuations by remodeling the abundance of plasma membrane (PM) transporters through regulated endocytosis (MacGurn et al., 2012; Léon and Teis, 2018). In budding yeast, the high-affinity methionine permease Mup1 provides a well-established model for this process. When methionine is limiting, Mup1 is stabilized at the PM to support methionine uptake, whereas methionine repletion or prolonged nutrient starvation trigger Mup1 internalization for vacuolar degradation, enabling rapid metabolic adaptation (Teis et al., 2008; Lin et al., 2008; Guiney et al., 2016; Ivashov et al., 2020; MacDonald et al., 2015). Despite extensive characterization of the endocytic pathway, the upstream cellular cues that determine how transporter remodeling is regulated remain incompletely understood.

Emerging evidence indicates that membrane lipid composition is a critical determinant of PM transporter behavior. Sphingolipids (SLs) are essential for PM organization and endocytic function (Zanolari et al., 2000; Wang and Chang, 2002; Finicle et al., 2018), and pharmacological inhibition of SL biosynthesis disrupts transporter activity and promotes selective internalization of Mup1 even under-nutrient replete conditions (Hepowit et al., 2021, 2023). This lipid-driven remodeling is sufficient to induce a cellular state resembling amino acid restriction and promotes stress resistance, highlighting SLs homeostasis as a major regulator of nutrient transporter function (Hepowit et al., 2021; Adebayo et al., 2025).

A genetic link between SL metabolism and Mup1 trafficking emerged from a genome-wide screen that identified the endoplasmic reticulum (ER)-vacuole tether Mdm1 as a factor required for proper Mup1 sorting (Henne et al., 2015). Loss of Mdm1 impaired Mup1 trafficking, and disease-associated Mdm1 truncations conferred hypersensitivity to the SL synthesis inhibitor myriocin, implicating this inter-organelle tether in SL regulation (Henne et al., 2015). Subsequent studies demonstrated that Mdm1 contributes to the intracellular flux of SL intermediates and coordinates lipid metabolic processes at the ER-vacuole contact sites (Girik et al., 2022; Hariri et al., 2018, 2019). Together, these observations position Mdm1 as a candidate organizer of lipid conditions that influence transporter fate at the PM.

However, whether Mdm1 directly links SL homeostasis to nutrient transporter remodeling and metabolic adaptation has not been established. Here, we show that Mdm1 is required to maintain SL balance necessary for starvation-induced clearance of Mup1. Loss of Mdm1 disrupts long-chain base (LCBs) and ceramide composition, impairs Mup1 internalization, and limits intracellular methionine accumulation. Restoring SL precursors rescues both Mup1 trafficking and metabolic defects, demonstrating that these phenotypes arise from altered lipid homeostasis. Finally, *mdm1*Δ cells exhibit enhanced stress resistance and extended chronological lifespan (CLS), consistent with a SL-dependent methionine-restricted state. These findings identify Mdm1 as a spatial regulator that couples lipid metabolism to Mup1 remodeling during nutrient stress.

## Results

### Mdm1 maintains sphingolipid homeostasis

To determine whether Mdm1 contributes to SL regulation, we performed targeted lipidomic profiling of key SL intermediates in wildtype and *mdm1*Δ cells grown under standard conditions. We quantified LCBs, including dihydrosphingosine (DHS) and phytosphingosine (PHS), as well as downstream ceramide species differing in acyl chain length (**Fig. 1A-C**).

**Figure 1:**
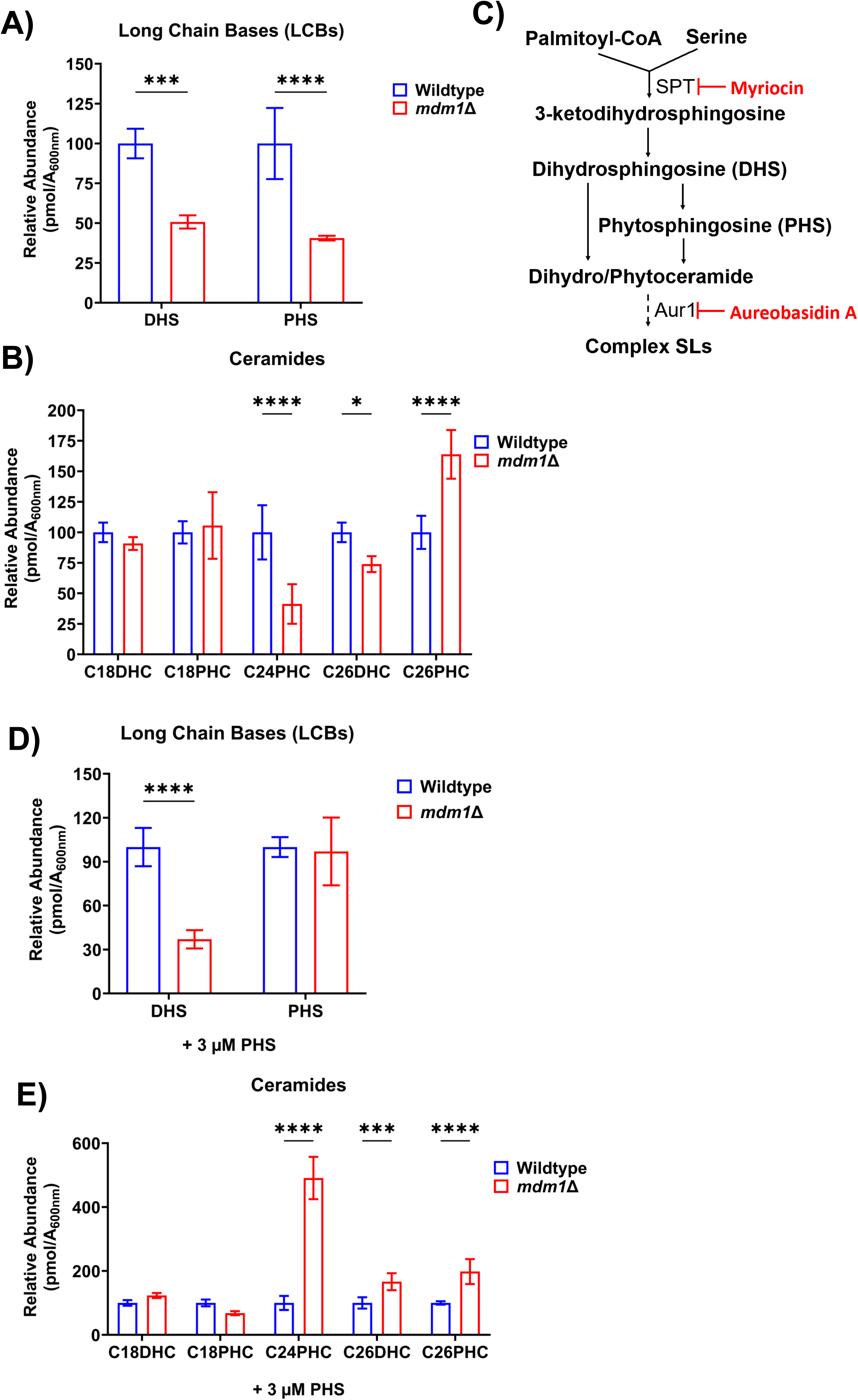
Mdm1 maintains sphingolipid homeostasis. A. Relative abundance of long-chain bases (DHS and PHS) in *mdm1*Δ relative to WT cells grown to log phase in SCD media (mean ± SD, n = 4, \*\*\**p* < 0.001, \*\*\*\**p* < 0.0001, Two-way ANOVA). B. Relative abundance of ceramide species in WT and *mdm1*Δ cells grown to log phase in SCD media (mean ± SD, n = 4, \**p* < 0.05, \*\*\*\**p* < 0.0001, Two-way ANOVA). C. Schematic of the sphingolipid biosynthetic pathway highlighting enzymatic steps inhibited by myriocin (serine palmitoyltransferase, SPT) and aureobasidin A (Aur1). D. LCB quantification in *mdm1*Δ relative to WT. Cells were grown in media containing 3 μM PHS (mean ± SD, n = 3, \**p* < 0.05, \*\*\*\**p* < 0.0001, Two-way ANOVA). E. Ceramide quantification in *mdm1*Δ relative to WT. Cells were grown in media containing 3 μM PHS (mean ± SD, n = 3, \*\*\**p* < 0.001, \*\*\*\**p* < 0.0001, Two-way ANOVA).

Loss of Mdm1 resulted in a pronounced reduction in total LCB abundance, with both DHS and PHS significantly decreased relative to wildtype cells (**Fig. 1A**). Consistent with reduced availability of free LCBs, levels of their phosphorylated derivatives, DHS-1P and PHS-1P, were also markedly diminished in *mdm1*Δ cells (**Fig. S1A**). Analysis of downstream ceramides revealed selective remodeling rather than a uniform reduction in all SL species. Specifically, very-long-chain ceramides C24PHC and C26DHC were reduced, whereas C26PHC accumulated to high levels, while shorter-chain ceramides (C18DHC and C18PHC) were largely unchanged (**Fig. 1B**). Together, these patterns indicate disruption of LCB availability and SL chain-length balance rather than a global suppression in SL synthesis in *mdm1*Δ cells. Notably, this altered SL landscape was observed in both logarithmic (log) and stationary growth phase conditions (**Fig. S1B**), suggesting that Mdm1 contributes broadly to SL homeostasis across growth states.

We next asked whether restoring LCB availability could bypass the defects in ceramide levels. Interestingly, exogenous supplementation with PHS, which circumvents early steps in *de novo* SL synthesis (Zanolari et al., 2000), restored cellular PHS levels in *mdm1*Δ cells to near-wildtype abundance, although DHS remained reduced (**Fig. 1C, D**). Downstream ceramides were partially normalized, including recovery of previously depleted C24PHC and C26DHC species, while elevated C26PHC persisted (**Fig. 1E**). Phosphorylated derivatives of DHS and PHS were also increased following PHS supplementation (**Fig. S1C**), indicating efficient uptake and metabolic utilization of exogenous LCBs.

Collectively, these findings demonstrate that Mdm1 is required to maintain SL homeostasis. Loss of Mdm1 remodels SL landscape by limiting LCB availability and subsequently disrupting ceramide balance, defects that could be rescued through PHS precursor supplementation.

### Loss of Mdm1 alters cellular adaptation to sphingolipid stress

Because disruption of SL homeostasis often influences cellular fitness under lipid stress, we next asked whether loss of Mdm1 alters growth response to pharmacological inhibition of the SL biosynthetic pathway. Prior work demonstrated that overexpression of disease-analogous truncation of Mdm1 induced hypersensitivity to myriocin, suggesting that perturbation of Mdm1 function disrupts SL homeostasis (Henne et al., 2015). However, whether loss of Mdm1 produces and similar outcome has not been examined.

We cultured wildtype and *mdm1*Δ cells in the presence of myriocin, an inhibitor of serine palmitoyltransferase (SPT) complex, which catalyzes the first step of *de novo* SL synthesis (**Fig. 1C**). During log growth, myriocin treatment reduced growth in wildtype cells by approximately 58% and in *mdm1*Δ cells by approximately 48% relative to vehicle-treated controls (**Fig. 2A; Fig. S2A**). Upon entry into stationary phase, myriocin continued to reduce growth in both wildtype and *mdm1*Δ cells; with ∼53% reduction in wildtype cells and ∼40% reduction in *mdm1*Δ cells (**Fig. 2B, Fig. S2A**). Therefore, myriocin treatment reduced growth in both strains; however, *mdm1*Δ cells exhibited a significant, albeit modest, attenuation in growth inhibition relative to wildtype across growth phases (**Fig. 2C)**. Thus, although *mdm1*Δ cells display reduced baseline fitness in response to myriocin treatment, their relative resistance to myriocin suggests an altered physiological response to SL limitation compared to wildtype cells.

**Figure 2:**
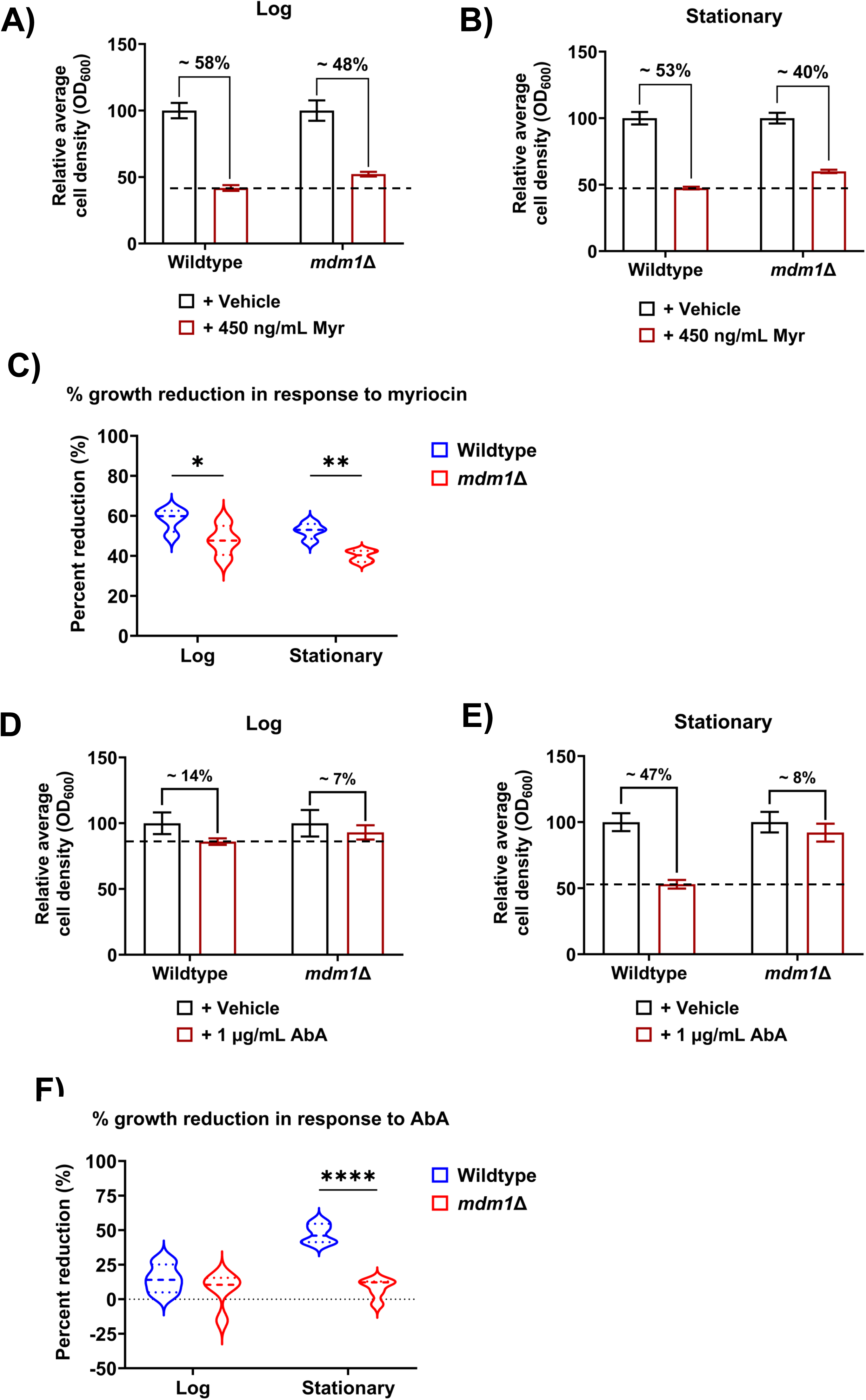
Loss of Mdm1 alters cellular adaptation to sphingolipid stress. A. Relative average cell density (OD₆₀₀) of wildtype and *mdm1*Δ cells grown in liquid culture with vehicle or 450 ng/mL myriocin (Myr) during log phase. Percent values indicate the magnitude of growth reduction relative to vehicle-treated controls (mean ± SD, n = 4). B. Relative average cell density (OD₆₀₀) of wildtype and *mdm1*Δ cells grown in liquid culture with vehicle or 450 ng/mL myriocin during stationary phase. Percent values indicate the magnitude of growth reduction relative to vehicle-treated controls (mean ± SD, n = 3). C. Quantification of percent growth reduction upon myriocin treatment in wildtype and *mdm1*Δ cells during log and stationary phases. Each point represents an independent experiment (mean ± SD; \**p* < 0.05, \*\**p* < 0.01; Two-way ANOVA). D. Relative average cell density (OD₆₀₀) of wildtype and *mdm1*Δ cells grown with vehicle or 1 µg/mL aureobasidin A (AbA) during log phase. Percent values indicate the magnitude of growth reduction relative to vehicle-treated controls (mean ± SD, n = 3). E. Relative average cell density (OD₆₀₀) of wildtype and *mdm1*Δ cells grown with vehicle or 1 µg/mL aureobasidin A (AbA) during stationary phase. Percent values indicate the magnitude of growth reduction relative to vehicle-treated controls (mean ± SD, n = 3). F. Quantification of percent growth reduction upon AbA treatment in wildtype and *mdm1*Δ cells during log and stationary phases. Each point represents an independent experiment (mean ± SD; \*\*\**p* < 0.0001; Two-way ANOVA.).

To determine whether this phenotype reflected a general response to SL perturbation rather than a myriocin-specific effect, we treated cells with aureobasidin A (AbA), an inhibitor of inositolphosphorylceramide synthase (Aur1) that targets a downstream step in SL synthesis (**Fig. 1C**). Consistent with the myriocin response, *mdm1*Δ cells exhibited attenuated growth inhibition compared to wildtype, particularly in stationary phase (**Fig. S2B**). During log growth, AbA caused a slight reduction in growth in wildtype cells (∼14%), whereas *mdm1*Δ cells were minimally affected (∼7%) relative to vehicle-treated controls (**Fig. 2D)**. Strikingly, in stationary phase, AbA treatment led to a pronounced reduction in wildtype growth (∼47%), while *mdm1*Δ cells again exhibited only a minor decrease (∼8%) (**Fig. 2E**). Consistent with this, growth resistance of *mdm1*Δ cells in response to AbA was observed in plating assays (**Fig. S2C**). Therefore, *mdm1*Δ cells exhibited a significant attenuation in growth inhibition in response to AbA treatment relative to wildtype particularly in stationary phase (**Fig. 2F)**.

Together, these findings indicate that loss of Mdm1 alters growth responses to SL biosynthetic stress, consistent with the lipidomic evidence of disrupted SL composition in *mdm1*Δ cells, suggesting that Mdm1 is required to maintain SL homeostasis. We therefore asked whether this SL imbalance directly contributes to methionine transporter Mup1 trafficking defects previously reported in *mdm1*Δ cells (Henne et al., 2015).

### Sphingolipid imbalance impairs starvation-induced clearance of Mup1

Previous work identified Mdm1 in a genome-wide screen for factors required for proper sorting for the methionine transporter Mup1 following methionine repletion, where loss of Mdm1 resulted in Mup1 trafficking defects (Henne et al., 2015). However, whether Mdm1 is also required for adaptive remodeling of Mup1 during prolonged starvation, and whether its effects on Mup1 trafficking are mediated through altered SL homeostasis, has remained unknown. Therefore, we directly tested the hypothesis that SL imbalance in *mdm1*Δ cells impairs starvation-induced clearance of Mup1 from the PM during starvation.

To monitor Mup1 trafficking, we used an endogenously tagged Mup1-pH reporter, in which pH-sensitive fluorophore is quenched upon delivery to the acidic vacuolar lumen but remains fluorescent at the PM or in early endocytic compartments (**Fig. 3A)** (Prosser et al., 2010; Ivashov et al., 2020; Henne et al., 2015). Wildtype cells expressing Mup1-pH were grown in methionine-free (-Met) and imaged during log and stationary phase (24hr). As expected, Mup1-pH localized predominantly to the PM during log growth in -Met, consistent with stabilization of Mup1 at the cell surface under methionine depletion to promote methionine uptake (**Fig. 3B)**. In contrast, in stationary phase, PM-localized Mup1-pH fluorescence was largely absent, consistent with starvation-induced endocytosis of Mup1 and delivery to the vacuole where Mup1-pH fluorescence was quenched (**Fig. 3B**).

**Figure 3:**
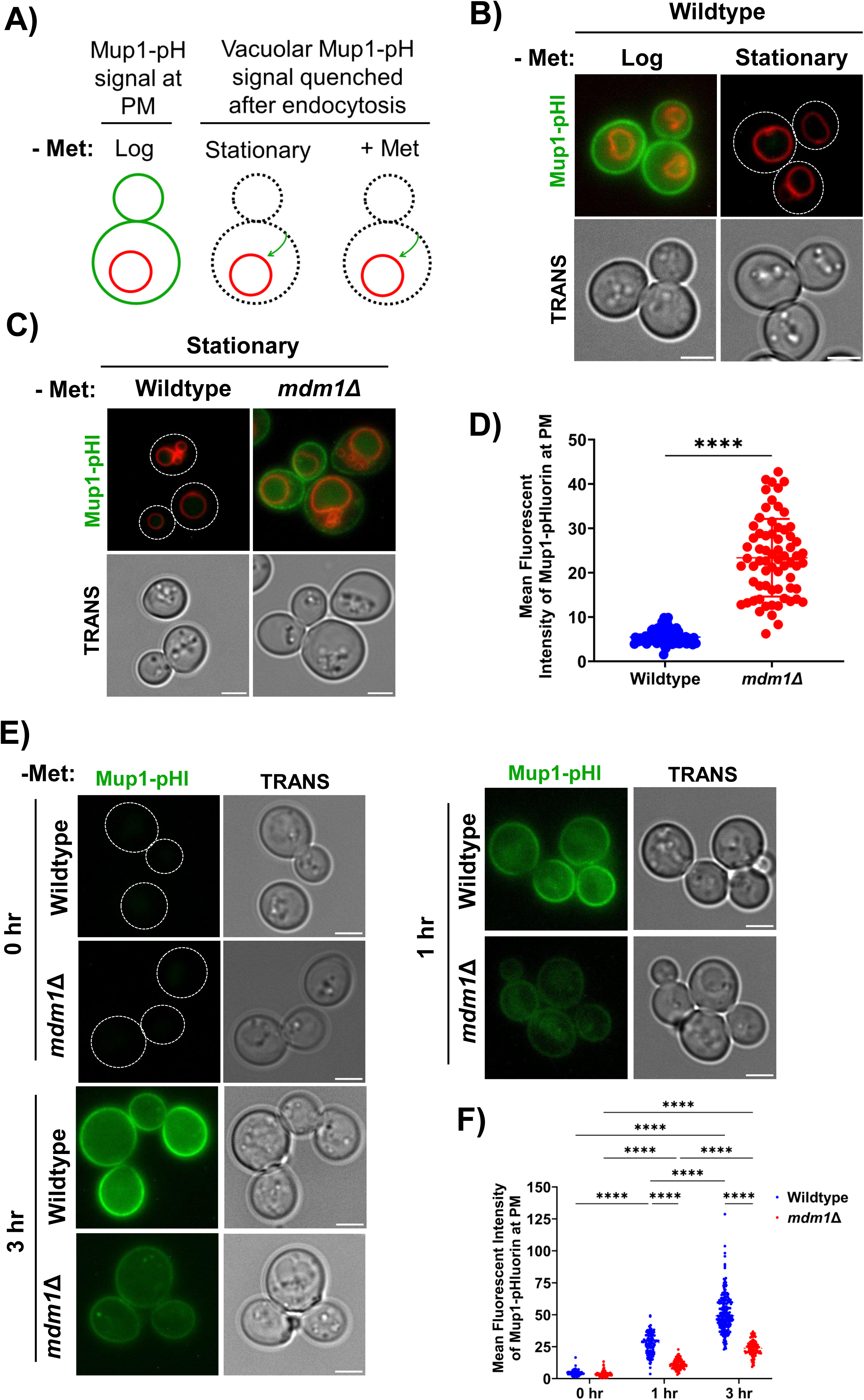
Sphingolipid imbalance impairs starvation-induced clearance of the methionine transporter Mup1. A. Schematic of the Mup1-pH assay. In methionine-free (−Met) conditions during log growth, Mup1-pH localizes to the plasma membrane (PM) and remains fluorescent. During stationary phase or after methionine re-addition (+Met), Mup1 undergoes endocytosis and delivery to the vacuole, where the pH-sensitive signal is quenched. B. Wildtype cells expressing endogenously tagged Mup1-pH with labeled vacuoles (FM4-64) grown in −Met and imaged during log growth or after prolonged growth to stationary phase. Dashed outlines indicate cell boundaries. TRANS images are shown below, Scale bar, 3 μM. C. Wildtype and *mdm1*Δ cells expressing endogenously tagged Mup1-pH with labeled vacuoles (FM4-64) grown in −Met and imaged at stationary phase. Dashed outlines indicate cell boundaries. TRANS images are shown below, Scale bar, 3 μM. D. Quantification of mean PM Mup1-pH fluorescence intensity from cells in **C**. Data represent individual cells (mean ± SD, n > 75 cells, \*\*\*\**p* < 0.0001, students *t*-test). E. Wildtype and *mdm1*Δ cells expressing Mup1-pH were shifted from +Met to −Met and imaged at 0, 1, and 3 hr to monitor synthesis and PM delivery of newly induced Mup1. TRANS images are shown below, Scale bar, 3 μM. F. Quantification of mean PM Mup1-pH fluorescence intensity from cells in **D**. Data represent individual cells (mean ± SD, n > 100 cells, \*\*\*\**p* < 0.0001, two-way ANOVA).

We next asked whether Mdm1 is required for this adaptive clearance. Wildtype and *mdm1*Δ cells expressing Mup1-pH were grown to stationary phase in -Met and analyzed by live-cell imaging. Whereas wildtype cells efficiently removed Mup1-pH from the PM under starvation conditions (SP), *mdm1*Δ cells retained a relatively stronger PM-localized Mup1-pH fluorescence signal (**Fig. 3C**). Quantification of this signal confirmed a significant increase in mean PM Mup1-pH intensity in *mdm1*Δ cells relative to wildtype, suggesting impaired starvation-induced Mup1-pH clearance in those cells (**Fig. 3D**).

Persistent PM localization of Mup1 could arise either from defective endocytic clearance or from increased synthesis and delivery of newly produced transporter to the cell surface. To distinguish between these possibilities, we examined the appearance of newly synthesized Mup1 at the PM using an established methionine depletion-induction assay (Hepowit et al., 2021). In this assay, wildtype and *mdm1*Δ cells expressing endogenously tagged Mup1-pH were shifted from methionine-replete to methionine-free medium to induce Mup1 expression and PM transport and monitored Mup1-pH signal over time.

As expected, in methionine-replete medium, both wildtype and *mdm1*Δ cells displayed no Mup1-pH signal at the PM (0 hr) (**Fig. 3E**). Within 1 hr of methionine depletion, wildtype cells displayed robust accumulation of Mup1-pH at the PM, consistent with efficient synthesis and trafficking, and continued to increase over time (3 hr). In contrast, *mdm1*Δ cells exhibited markedly reduced PM-localized Mup1-pH signal at early time points (1 hr) and a slight increase at 3 hr time points, indicating impaired delivery of newly synthesized transporter to the cell surface (**Fig. 3E, F)**. These findings argue against increased Mup1 synthesis as the basis for the persistent PM localization observed in *mdm1*Δ cells, and instead, are more consistent with impaired endocytic clearance of Mup1 in *mdm1*Δ cells (**Fig. 3C**). Supporting this this interpretation, immunoblot analysis of Mup1-pH during methionine depletion revealed reduced steady-state abundance in *mdm1*Δ cells (**Fig. S3A, B)**.

Together, these findings demonstrate that Mdm1 is required for efficient starvation-induced remodeling of Mup1. Although overall Mup1 abundance is reduced, Mup1 fails to be properly cleared from the PM in *mdm1*Δ cells, consistent with a defect in adaptive endocytic turnover.

### Loss of Mdm1 limits intracellular methionine accumulation

Because Mup1 is the primary high-affinity methionine transporter in budding yeast, we next asked whether impaired starvation-induced clearance of Mup1 in *mdm1*Δ cells is associated with the altered intracellular methionine availability. We therefore performed targeted metabolomic analysis on wildtype and *mdm1*Δ cells grown in synthetic complete medium (SCD) to assess steady-state nutrient levels. Under these conditions, *mdm1*Δ cells exhibited a substantial reduction in intracellular methionine relative to wildtype cells (**Fig. 4A**), indicating that loss of Mdm1 limits methionine accumulation in cells even when extracellular methionine is available.

**Figure 4:**
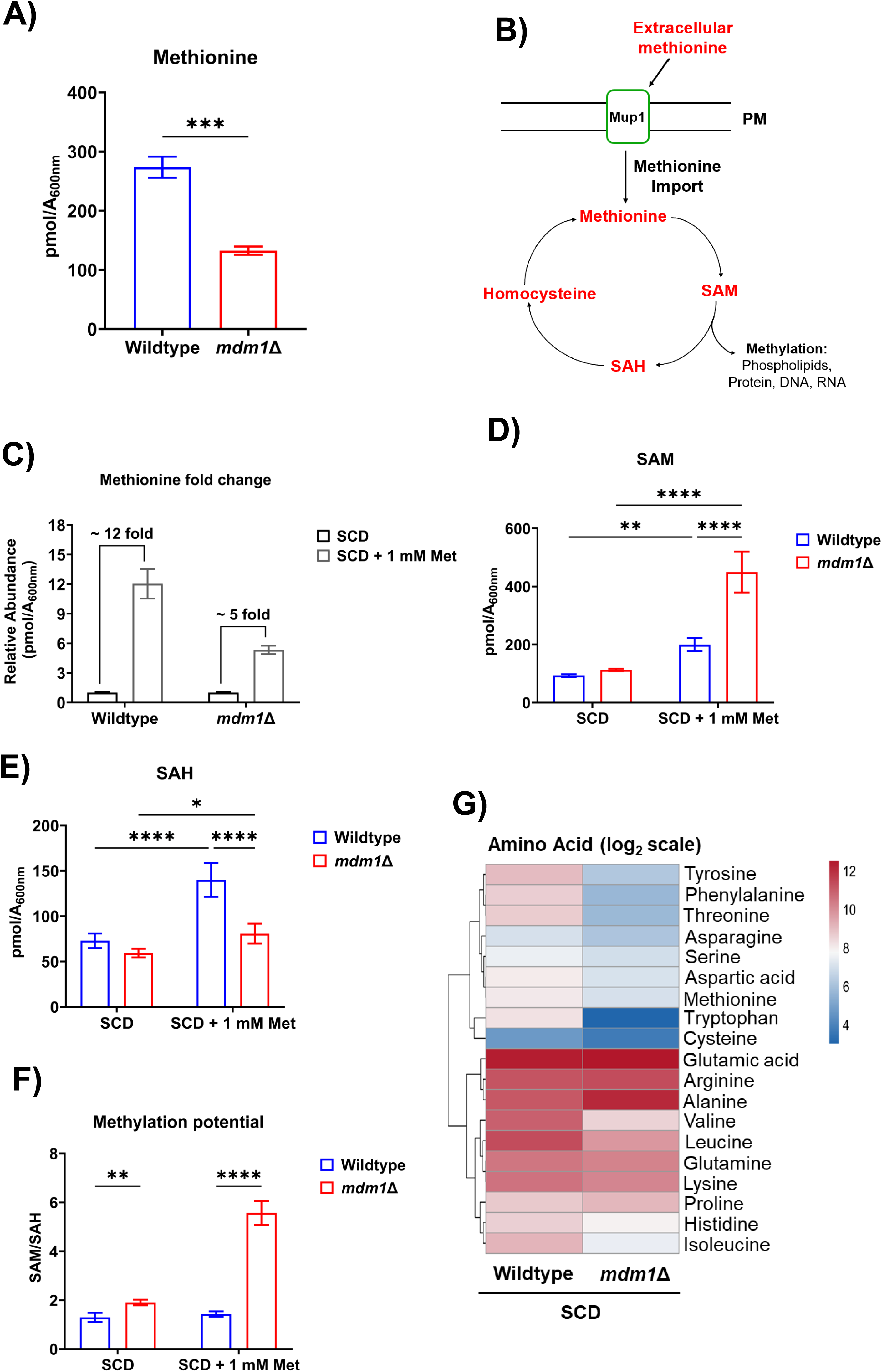
Loss of Mdm1 limits intracellular methionine accumulation. A. Relative change in intracellular methionine abundance in wildtype and *mdm1*Δ cells upon supplementation with 1 mM methionine. Values are normalized to SCD conditions. Bars represent mean ± SD, n = 3, \**p* < 0.05, students *t*-test. B. Schematic illustrating Mup1-mediated methionine import and its incorporation into the methionine cycle, showing conversion between methionine, S-adenosylmethionine (SAM), S-adenosylhomocysteine (SAH), and homocysteine (Hcy). C. Relative fold change in intracellular methionine abundance in wild-type and *mdm1*Δ cells upon supplementation with 1 mM methionine. Values are normalized to SCD conditions. Bars represent mean ± SD, n = 3, \**p* < 0.05, students *t*-test. D. Intracellular SAM abundance in wildtype and *mdm1*Δ cells grown in SCD or SCD supplemented with 1 mM methionine. Bars represent mean ± SD, n = 3 biological replicates, \*\**p* < 0.01, \*\*\*\**p* < 0.0001, Two-way ANOVA. E. Quantification of SAH in wildtype and *mdm1*Δ cells grown in SCD or SCD supplemented with 1 mM methionine (mean ± SD, n = 3, \**p* < 0.05, \*\*\*\**p* < 0.0001, Two-way ANOVA). F. Methylation potential, expressed as the SAM:SAH ratio, in wildtype and *mdm1*Δ cells grown in SCD or SCD + 1 mM methionine. Bars represent mean ± SD, n = 3 biological replicates, \*\**p* < 0.01, \*\*\*\**p* < 0.0001, Two-way ANOVA. G. Heatmap showing log₂ fold-change in intracellular amino acid abundance (wildtype relative to *mdm1*Δ) in SCD. Color scale represents log₂ fold-change.

Reduced intracellular methionine could arise either from impaired extracellular uptake or from increased downstream utilization (**Fig. 4B**). To distinguish between these possibilities, we first supplemented cultures with excess methionine and quantified intracellular levels. In wildtype cells, methionine supplementation resulted in a ∼12-fold increase in intracellular methionine levels (**Fig. 4C**). In contrast, *mdm1*Δ cells showed a blunted increase (∼5-fold increase), and absolute methionine levels remained substantially lower compared to wildtype cells (**Fig. S4A**). These results indicate that *mdm1*Δ cells exhibit a diminished capacity to accumulate methionine even when it is abundant in the environment.

We next examined intermediates of the methionine cycle to evaluate whether increased metabolic consumption could account for the methionine reduction observed in *mdm1*Δ cells. Methionine is converted to S-adenosylmethionine (SAM), which serves as a universal methyl donor in diverse methylation reactions (**Fig. 4B**). Following methyl transfer, SAM is converted to S-adenosylhomocysteine (SAH), which is subsequently hydrolyzed to homocysteine. Thus, SAH and homocysteine abundance provide indirect readout of methylation flux through the cycle.

Despite reduced intracellular methionine, *mdm1*Δ cells displayed elevated SAM and reduced SAH upon addition of exogenous methionine, resulting in a significantly increased SAM:SAH ratio relative to wildtype cells (**Fig. 4B, D, E**). Homocysteine levels were also reduced (**Fig. S4B)**. Because both SAH and homocysteine are generated downstream of SAM-dependent methylation, their reduced abundance argues against increased methionine flux through the cycle and instead supports a model in which intracellular methionine is limited by impaired uptake.

Consistent with this interpretation, global amino acid profiling revealed that, in addition to methionine, multiple intracellular amino acids were also reduced in *mdm1*Δ cells (**Fig. 4G; Fig. S4C**). This included serine, which is a key metabolite that contributes to SL biosynthesis. However, some amino acids were either unchanged or elevated, suggesting that the metabolic impact of loss of Mdm1 reflects selective remodeling of amino acid pools. While we did not directly measure uptake of additional transporters, the overall pattern is consistent with altered transporter capacity, thereby linking Mdm1-mediated SL imbalance to nutrient availability.

### Restoration of sphingolipid precursors rescues transporter and metabolic defects

Our lipidomic analysis revealed a marked reduction in LCBs in *mdm1*Δ cells, raising the possibility that impaired SL availability underlies the observed defects in Mup1 remodeling and amino acid levels (**Fig. 1A)**. We therefore asked whether replenishing LCBs is sufficient to restore endocytic remodeling.

To directly test this model, we supplemented wildtype and *mdm1*Δ cultures with 3 µM PHS and examined starvation-induced Mup1 trafficking using the stationary phase Mup1-pH reporter (**Fig. 3A)**. Strikingly, PHS supplementation markedly enhanced clearance of PM-localized Mup1 in *mdm1*Δ cells (**Fig. 5A)**, and quantification confirmed a significant reduction in PM fluorescent intensity (**Fig. 5B)**, indicating that restoring SL precursors is sufficient to rescue starvation-induced Mup1 clearance in *mdm1*Δ cells.

**Figure 5:**
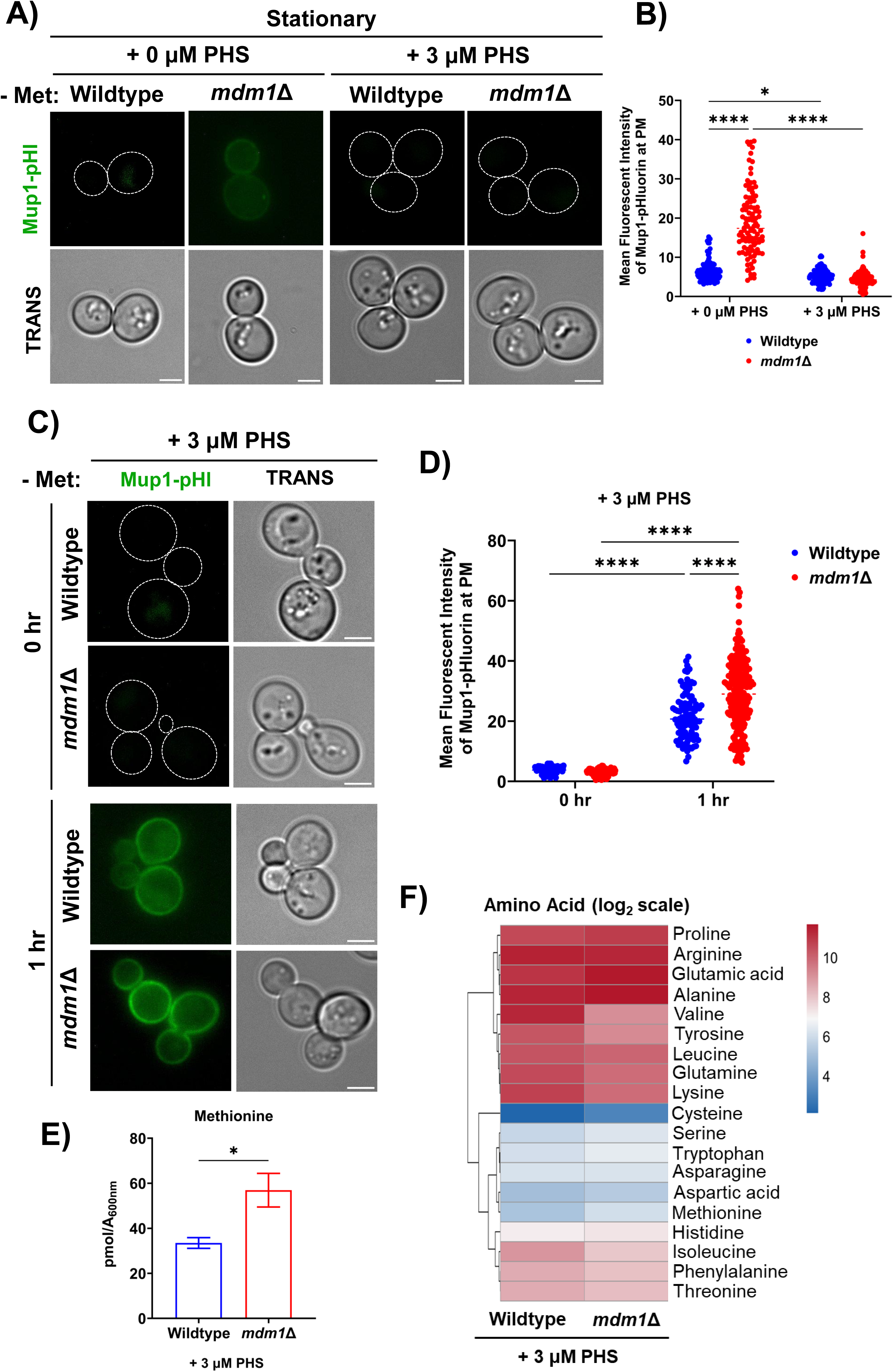
Restoration of sphingolipid precursors rescues transporter and metabolic defects. A. Wildtype and *mdm1*Δ cells expressing endogenously tagged Mup1-pH grown in −Met containing 0 μM or 3 μM PHS and imaged at stationary phase. Dashed outlines indicate cell boundaries. TRANS images are shown below, Scale bar, 3 μM. B. Quantification of mean PM Mup1-pH fluorescence intensity from cells in **A**. Data represent individual cells (mean ± SD, n > 100 cells, \*\**p* < 0.01, students *t*-test). C. Wildtype and *mdm1*Δ cells expressing Mup1-pH were shifted from +Met to −Met containing 3 μM PHS and imaged at 0 and 1 hr to monitor synthesis and PM delivery of newly induced Mup1. TRANS images are shown below, Scale bar, 3 μM. D. Quantification of mean PM Mup1-pH fluorescence intensity from cells in **C**. Each point represents an individual cell (mean ± SD, n > 100 cells, \*\*\*\**p* < 0.0001, students *t*-test). E. Intracellular methionine abundance in wildtype and *mdm1*Δ cells grown in SCD medium containing 3 μM PHS. Bars represent mean ± SD, n = 3, \**p* < 0.05, students *t*-test. F. Heatmap showing log₂ fold-change in intracellular amino acid abundance (wildtype relative to *mdm1*Δ) following 3 μM PHS supplementation. Color scale represents log₂ fold-change.

To determine whether this rescue reflected improved trafficking rather than reduced transporter synthesis, we examined Mup1 PM delivery following methionine depletion in wildtype and *mdm1*Δ cells. Importantly, PHS supplementation enhanced Mup1 PM localization in *mdm1*Δ cells (**Fig. 5C, D**), demonstrating recovery of adaptive trafficking dynamics.

Finally, we examined whether restoration of Mup1 remodeling translates into metabolic recovery. Indeed, targeted metabolomics analysis revealed that PHS supplementation significantly increased intracellular methionine levels in *mdm1*Δ cells (**Fig. 5E**). Global amino acid profiling similarly showed broad attenuation of the amino acid depletion phenotype, although rescue remained incomplete for some metabolites suggesting that while LCB replenishment restores a major component of transporter function, Mdm1 loss likely causes broader metabolic defects that cannot be fully bypassed by exogenous PHS (**Fig. 5F; Fig. S5D**). Consistent with improved methionine availability, intermediates of the methionine cycle exhibited partial normalization, including reduction of the elevated SAM:SAH ratio in *mdm1*Δ (**Fig. S5A–C**).

Together, these findings demonstrate that SL availability is a key determinant of adaptive transporter remodeling and nutrient accumulation. Restoring LCB levels is sufficient to rescue both trafficking and metabolic defects in *mdm1*Δ cells, establishing altered SL homeostasis as a primary driver of the mutant phenotype.

### Loss of Mdm1 promotes stress resistance and extends chronological lifespan

The SL-dependent defects in Mup1 remodeling observed in *mdm1*Δ cells result in persistently reduced intracellular methionine and broader amino acid depletion, a metabolic state reminiscent of nutrient restriction. Because both methionine limitation and reduced SL synthesis have been associated with enhanced stress resistance and longevity, we asked whether loss of Mdm1 influences long-term cellular survival (Johnson and Johnson, 2014; Zou et al., 2020; Huang et al., 2012).

To test this, we performed chronological lifespan (CLS) assays, which measure the survival of non-dividing cells maintained in stationary phase. Interestingly, *mdm1*Δ cells exhibited a significant extension of CLS compared to wildtype cells (**Fig. 6A**), indicating enhanced survival under prolonged nutrient limitation. Notably, *MDM1* transcript levels declined during aging in wildtype yeast (**Fig. 6B**), suggesting that reduced Mdm1 expression may accompany physiological adaptation to long-term nutrient stress.

**Figure 6:**
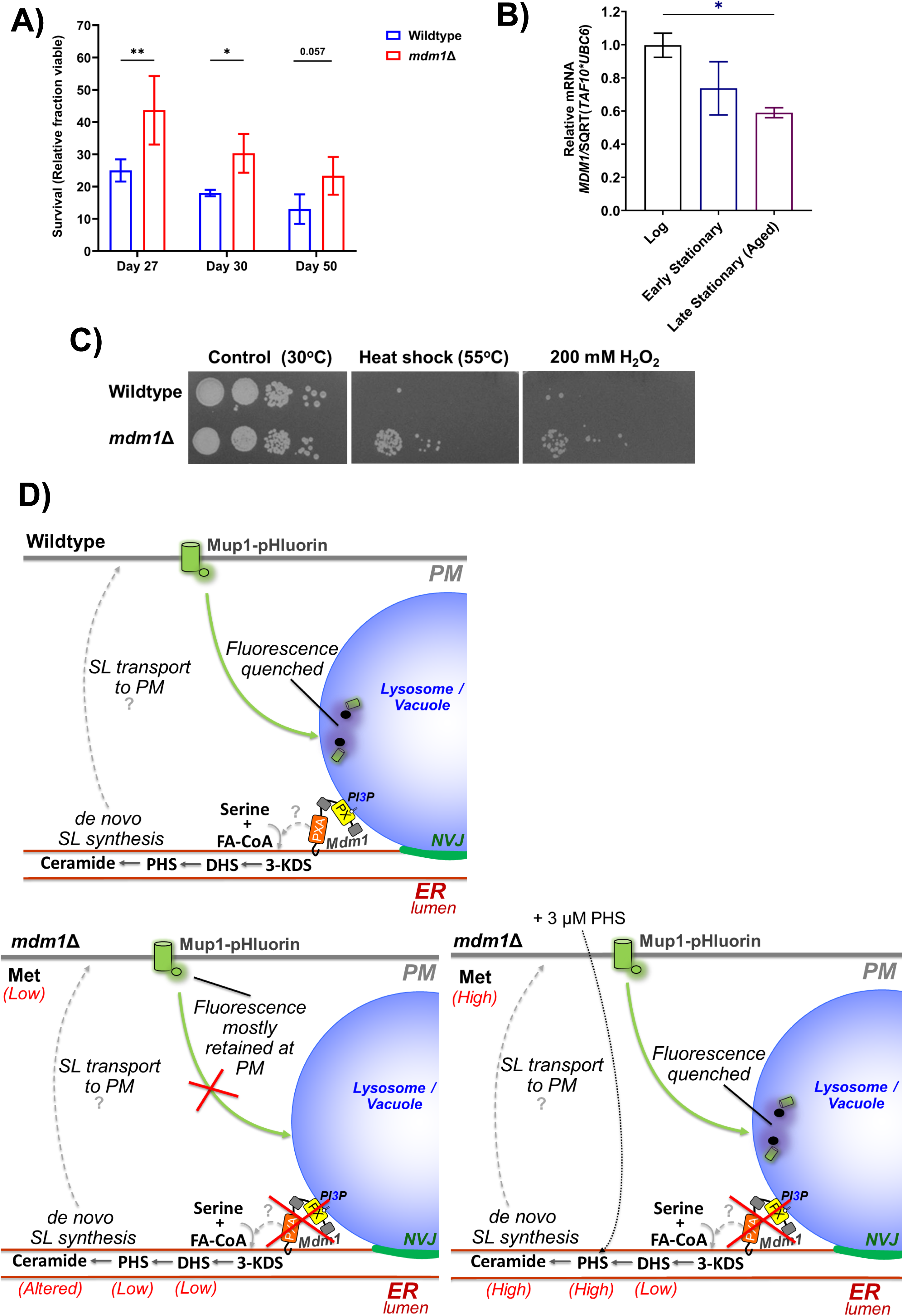
Loss of Mdm1 promotes stress resistance and extends chronological lifespan. A. CLS of WT and *mdm1*Δ cells. Statistical significance is indicated for specific time points (mean ± SD, n = 3 biological replicates, \**p* < 0.05, \*\*\**p* < 0.001, Two-way ANOVA). B. Resistance of *mdm1*Δ cells to heat (55 °C) or hydrogen peroxide (H_2_O_2_) stress. Photographs show a ten-fold dilution series of cells (from left to right). C. Relative expression of *MDM1* mRNA measured by qRT-PCR in wildtype cells during logarithmic growth, early stationary phase, and late stationary (aged) phase. Transcript levels were normalized to *TAF10* and *UBC6* (mean ± SEM, n = 3 biological replicates, \**p* < 0.05, One-way ANOVA). D. Model illustrating how Mdm1 couples sphingolipid (SL) metabolism to plasma membrane (PM) remodeling of the methionine transporter Mup1. **Top**, in wildtype cells, Mdm1 maintains SL homeostasis, enabling efficient endocytic trafficking of Mup1-pHluorin to the vacuole where fluorescence is quenched. **Bottom left**, loss of Mdm1 disrupts de novo SL balance (reduced long-chain bases and altered ceramides), causing defective Mup1 clearance, persistent PM retention, and reduced intracellular methionine. **Bottom right**, PHS supplementation restores SL pools in *mdm1*Δ cells, rescuing Mup1 endocytosis and increasing intracellular methionine. KDS, 3-ketodihydrosphingosine; DHS, dihydrosphingosine; PHS, phytosphingosine.

Finally, we examined whether *mdm1*Δ cells display enhanced resistance to acute stress. Following heat shock (55 °C) and oxidative stress (200 mM H_2_O_2_), *mdm1*Δ cells showed increased survival relative to wildtype populations (**Fig. 6C**), consistent with a stress-resistant cellular state.

Together, these findings identify loss of Mdm1 as a determinant of cellular adaptation to nutrient limitation. By perturbing SL homeostasis and altering Mup1-dependent methionine availability, *mdm1*Δ cells adopt a metabolic configuration associated with improved stress tolerance and extended lifespan.

## Discussion

Endocytic regulation of PM transporters is a central mechanism by which eukaryotic cells coordinate nutrient availability with metabolic state (MacGurn et al., 2012; Léon and Teis, 2018). In this study, we identify the NVJ protein Mdm1 as a regulator of SL homeostasis that coordinates Mup1 dynamics, metabolic balance, and cellular aging. Our data support a model in which Mdm1 maintains a lipid environment that is permissive for proper Mup1 endocytic recycling (**Fig. 6D**). In the absence of Mdm1, altered ceramides composition disrupts starvation-induced Mup1 clearance, limits intracellular methionine abundance, and promotes stress-resistant, longevity-associated metabolic state.

Previous work identified Mdm1 in a genome-wide screen for factors required for methionine-induced Mup1 sorting, where *mdm1*Δ cells exhibited trafficking defects (Henne et al., 2015). Those assays probed acute, signal-triggered internalization following methionine repletion (Henne et al., 2015). In contrast, our study examined Mup1 dynamics during prolonged starvation in stationary phase, a physiologically distinct condition that requires sustained endocytic turnover rather than transient signaling. Under these conditions, wildtype cells efficiently cleared Mup1 from the PM, whereas *mdm1*Δ cells retained the transporter at the cell surface despite reduced overall steady-state Mup1 abundance (**Fig. 3C – F**). These findings suggest that *mdm1*Δ cells are not defective in triggering endocytosis per se but are impaired in maintaining adaptive transporter clearance during chronic nutrient stress. Our findings therefore extend prior work by revealing a growth-phase-specific requirement for Mdm1 in sustained remodeling of Mup1.

Our data indicate that this trafficking defect arise from disrupted SL homeostasis rather than altered Mup1 synthesis. Lipidomic analysis revealed reduced LCBs and imbalanced ceramide composition in *mdm1*Δ cells (**Fig. 1A, B**), consistent with prior evidence that Mdm1 contributes to SL metabolism at ER-vacuole contact sites (Henne et al., 2015; Girik et al., 2022). Importantly, supplementation with PHS restored LCB levels, recued starvation-induced Mup1 clearance, and normalized intracellular methionine pools (**Fig. 1D; Fig. S1C; Fig. 5A, B, E**). These findings suggest that Mdm1 maintains SL homeostasis necessary for selective endocytic remodeling.

The requirement for SL homeostasis in transporter remodeling is consistent with accumulating evidence that SL depletion selectively alters PM proteostasis (Hepowit et al., 2021, 2023). Pharmacological inhibition of SL synthesis triggers Art2-dependent clearance of Mup1 and triggers a state resembling amino acid restriction (Hepowit et al., 2023). Our findings complement this model by demonstrating that endogenous disruption of SL balance through loss of Mdm1 similarly compromises Mup1 regulation even in the absence of pharmacological inhibition. Notably, *mdm1*Δ cells exhibit attenuated sensitivity to inhibitors of SL biosynthesis, suggesting chronic adaptation to altered lipid flux (**Fig. 2; Fig. S2**). Together, these observations support a model in which Mdm1 maintains SL precursor availability and ceramide balance required for proper PM organization and endocytic remodeling (**Fig. 1**).

The trafficking defects observed in *mdm1*Δ cells have direct metabolic consequences. Despite extracellular methionine availability, *mdm1*Δ cells display reduced intracellular methionine levels and a broader decrease in amino acid pools (**Fig. 4C, G**). Metabolite profiling argues against increased methionine consumption and instead support impaired Mup1-mediated accumulation (**Fig. 4D, E**). These findings align with the emerging view that SL metabolism and methionine homeostasis are functionally linked (Adebayo et al., 2025; Hepowit et al., 2021, 2023). In this model, disruption of SL composition alters nutrient transporter function, thereby restricting intracellular amino acid availability and downstream one-carbon metabolism. Therefore, Mdm1 links lipid metabolic state to nutrient sensing and metabolic programming.

The metabolic state induced by Mdm1 loss closely resembles methionine restriction, a widely observed longevity-promoting intervention across eukaryotes (Lee et al., 2014; Kozieł et al., 2014; Plummer and Johnson, 2019) suggesting that disruption of SL-dependent transporter regulation phenocopies nutrient-restricted aging programs. *mdm1*Δ cells exhibit enhanced resistance to heat and oxidative stress and extended chronological lifespan (**Fig. 6C**). Reduced expression of *MDM1* during chronological aging further suggests that modulation of Mdm1 activity may contribute to age-associated metabolic remodeling (**Fig. 6B**). While we do not establish causality between Mdm1-mediated SL homeostasis and longevity, our findings support a model in which chronic, SL disruption promotes a nutrient state that enhances stress adaptation.

More broadly, our work highlights membrane contact sites as regulatory hubs that couple lipid metabolism to PM proteostasis. Mdm1 localizes to ER-vacuole contact sites and has been implicated in coordinating lipid flux and organelle lipid storage (Hariri et al., 2018, 2019). Our findings extend this role by demonstrating that SL imbalance at MCSs influences nutrient transporter turnover at the PM (**Fig. 6D**). Therefore, we propose that Mdm1 maintains a lipid environment permissive for adaptive endocytic remodeling, thereby integrating inter-organelle lipid metabolism with nutrient uptake and metabolic state. Loss of Mdm1 perturbs this lipid environment, impairing both induction and clearance of transporters such as Mup1, limiting methionine uptake, and triggering a nutrient-restricted metabolic state that promotes stress resistance and longevity. Defining whether Mdm1 primarily regulates SL flux, or compartmentalization at the ER-vacuole interface will be an important next step in understanding how spatial lipid regulation shapes cellular physiology.

## Materials and methods

### Molecular biology, yeast genetics, and growth conditions

Yeast genetic manipulations were conducted using classical yeast knock-in/out protocols. The strains used are listed in Tables S1. Yeast transformations were performed using the lithium acetate method. Strains were selected using antibiotics. All chemicals used to make the yeast media were purchased from Sigma-Aldrich (succinic acid, sodium hydroxide, ammonium sulfate, yeast nitrogen base without amino acids, or ammonium sulfate). Yeast media was supplemented with a final concentration of 2% dextrose. Phytosphingosine was purchased from Cayman chemical and 3 µM was used in the study. Myriocin and aureobasidin A were also purchased from Cayman chemical and concentrations used were 450 ng/mL and 1 µg/mL respectively.

### Growth assays

Precultures were grown in Synthetic Complete medium (SCD) containing 2% dextrose at 30 °C with orbital shaking (225 rpm). For growth assays, cultures were diluted to an OD₆₀₀ of 0.05 in their respective media and dispensed (200 µL per well) into 96-well plates with at least four technical replicates per biological sample. Plates were lidded, incubated at 30 °C in a SpectraMax i3x microplate reader with continuous shaking, and OD₆₀₀ readings were taken every 20 minutes. Raw absorbance values were blank-corrected by subtracting medium-only wells. Growth curves were plotted in GraphPad Prism, and average cell density were calculated by averaging the total density at log or stationary phases. All data are presented as mean ± SD from a minimum of four technical replicates from an independent experiment.

### Yeast spotting assay

Yeast strains were cultured under appropriate conditions, then serially diluted 10-fold in 96-well plates. Dilution series were replica-pinned onto YPD agar plates and incubated at 30 °C. Plates were imaged after 2 days to assess growth phenotypes.

### qPCR

Yeast strains were cultured under appropriate conditions, pelleted by centrifugation (3,000 × g, 5 min), and stored at −80 °C for RNA extraction. Upon thawing on ice, total RNA was extracted using the Takara kit per the manufacturer’s instructions. Briefly, pellets were resuspended in 350 µL LBP buffer with acid-washed glass beads (≈0.5 vol) and lysed by bead beating (4 × 1 min cycles with 1 min ice rests). The crude lysate was clarified and RNA further purified. First-strand cDNA was synthesized with the PrimeScript RT reagent kit with genomic DNA eliminator (Takara Bio, RR047A). qPCR was performed in 10 µL reactions containing PowerUp SYBR Green Master Mix (Thermo Fisher, A25777) on a real-time instrument; expression levels were normalized to *UBC6* or *TAF10*. Primer efficiencies (95–105%) were empirically validated, and all primer sequences are listed in Table 1.

### SL analysis by MS

SL analysis was performed largely as described by Bielawski et al. (2006) (Bielawski et al., 2006). Briefly, yeast cultures were grown under the indicated conditions, harvested (3,000 × g, 5 min, 4 °C), washed twice, and wet pellet weights recorded. Samples were spiked with endogenous calibration standards and synthetic internal standards prior to lipid extraction using a single-phase CHCl₃:MeOH:H₂O (2:1:0.8, v/v/v) system with bead beating. Following phase separation and re-extraction, combined organic fractions were dried under N₂ and reconstituted in MeOH:H₂O (9:1, v/v) containing 5 mM ammonium formate. Lipids were separated by reverse-phase HPLC and analyzed on a triple-quadrupole mass spectrometer operating in positive-ion multiple reaction monitoring (MRM) mode. SL species were quantified using analyte-to–internal standard peak area ratios referenced to calibration curves. SL abundances were normalized to OD₆₀₀.

### Targeted metabolomic analysis by LC-MS/MS

Targeted metabolomic profiling was performed using an LC–MS/MS platform adapted from Bao et al (2019) (Bao et al., 2019). Yeast metabolites were extracted by cold methanol precipitation with minor modifications. Briefly, cell pellets were extracted with prechilled methanol (−80 °C), lysed, and centrifuged at 10,000 rpm for 10 min at 4 °C. The supernatant was collected, and the pellet was re-extracted with 80% methanol. Combined extracts were dried in a refrigerated centrifugal concentrator and reconstituted in acetonitrile:water (50:50, v/v). Metabolites were quantified on an AB SCIEX QTRAP 6500 LC–MS/MS system coupled to a Shimadzu Nexera UHPLC. Data acquisition was performed using Analyst 1.6 and processed with MultiQuant 3.0. To maximize metabolite coverage, samples were analyzed using three chromatographic modes: reverse-phase separation on a Synergi Polar-RP column, HILIC separation on an Atlantis HILIC Silica column, and ion-pair chromatography on an Atlantis T3 column. Eluents were monitored in both positive and negative ionization modes using multiple reaction monitoring (MRM). Instrument parameters were optimized using direct infusion of metabolite standards. Calibration curves (10 nM–10 µM) and quality control samples were included for quantification. Metabolite abundances were normalized to OD₆₀₀.

### Chronological lifespan assay (CLS)

Lifespan assays CLS was measured as previously described (Wei et al., 2008). Overnight yeast cultures grown at 30 °C in SCD medium were diluted into 10 mL fresh medium (in 50 mL conical tubes) to an initial OD₆₀₀ of 0.3. Cultures were incubated at 30 °C with orbital shaking (225 rpm). Viability after the specified number of days was assessed by serial dilution and plating on YPD agar. Following 2–3 days of incubation at 30 °C, colonies were counted, and viability was expressed as a percent fraction. Statistical significance was determined using a Two-way ANOVA.

### Western blot analysis

Yeast proteins were extracted using an alkaline lysis protocol adapted from Kushnirov (2000) (Kushnirov, 2000). Briefly, cell pellets were resuspended in 100 μL distilled water followed by 100 μL of 0.2 M NaOH, vortexed, and incubated at room temperature for 5 min. Cells were pelleted by centrifugation at 18,000 × g for 30 s and the supernatant was removed. The resulting pellet was resuspended in a volume of 1× Laemmli sample buffer proportional to the optical density (OD₆₀₀) of the culture (e.g., OD₆₀₀ = 4 was resuspended in 40 μL buffer), ensuring normalized protein loading across samples. Lysates were boiled at 95 °C for 5 min to denature proteins and centrifuged at 18,000 × g for 1 min to pellet insoluble debris. For SDS–PAGE, 10 μL of the clarified supernatant was loaded per lane. Proteins were transferred to PVDF membranes using a semi-dry transfer system. Membranes were blocked in 5% non-fat dry milk in TBST (20 mM Tris-HCl pH 7.5, 150 mM NaCl, 0.1% Tween-20) and incubated with mouse anti-GFP antibody (1:1,000; Roche or equivalent), followed by HRP-conjugated secondary antibodies. Signal was detected using enhanced chemiluminescence (ECL) reagents and imaged using the iBright FL1500 Imaging System (Thermo Fisher Scientific). Total protein stain was used for normalization and served as the loading control.

### Image analysis

Fluorescence intensity of Mup1-pH at the plasma membrane was measured using Fiji (ImageJ). Regions corresponding to the PM were manually traced using the freehand or polygon tool, and mean fluorescence intensities were extracted. Background signal was subtracted from each measurement.

## Acknowledgements

We thank the Stony Brook Cancer Center Biological Mass Spectrometry Shared Resource for expert assistance with Lipidomics analysis. We thank Miriam Greenberg laboratory members for sharing supplies. H. Hariri is supported by the National Institutes of Health National Institute of General Medical Sciences (R35GM150892), Wayne State University start-up funds and University Research Award, Karmanos Cancer Institute - Strategic Research Initiative Grant (KCI-SRIG), Richard Barber Interdisciplinary Research Program. The Microscopy, Imaging and Cytometry Resources Core is supported in part by NIH Karmanos Cancer Institute Center Grant (P30 CA22453) and Wayne State University. The Pharmacology and Metabolomics Core was supported in part by NIH Center grant P30 CA22453 to the Barbara Ann Karmanos Cancer Institute.

The authors declare no competing financial interest.

## Author contributions

D. Adebayo and H. Hariri, conception and design of the study, data interpretation, drafting and revising the manuscript. D. Adebayo, E. Obaseki, K. Vasudeva, M. Aboumourad, S. Miller, growth and spotting assay, imaging, image quantification, immunoblot analysis, data analysis, figure preparation, critical reading of the manuscript, generating yeast strains, technical and editorial assistance. A. Ostermeyer-Fay and D. Canals, sphingolipid analysis, X. Bao and J. Li, metabolomics.

## Supplemental material

**Table S1:**
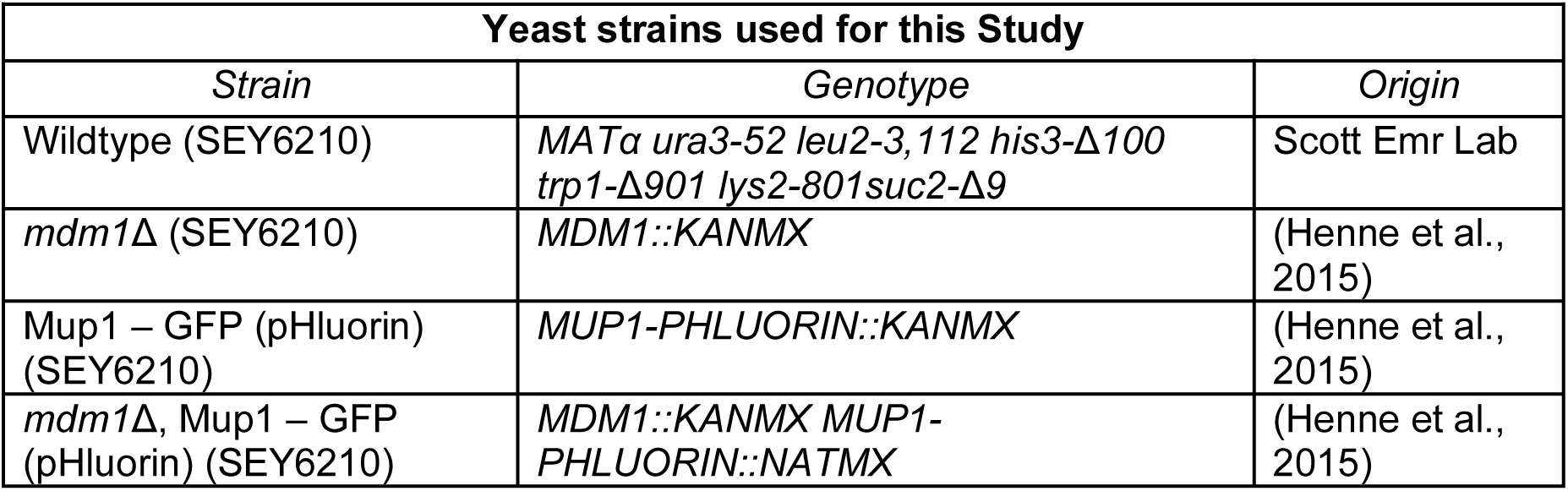
Yeast strains used in this study.

**Table S2:**
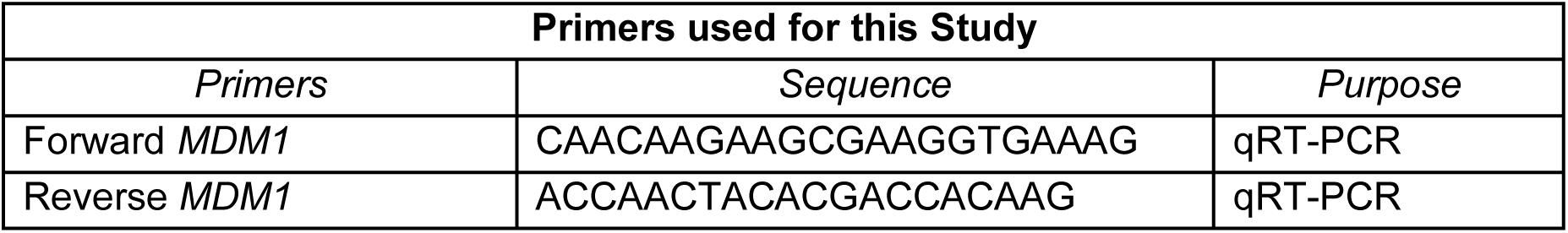
Primers used in this study.

**Figure S1.**
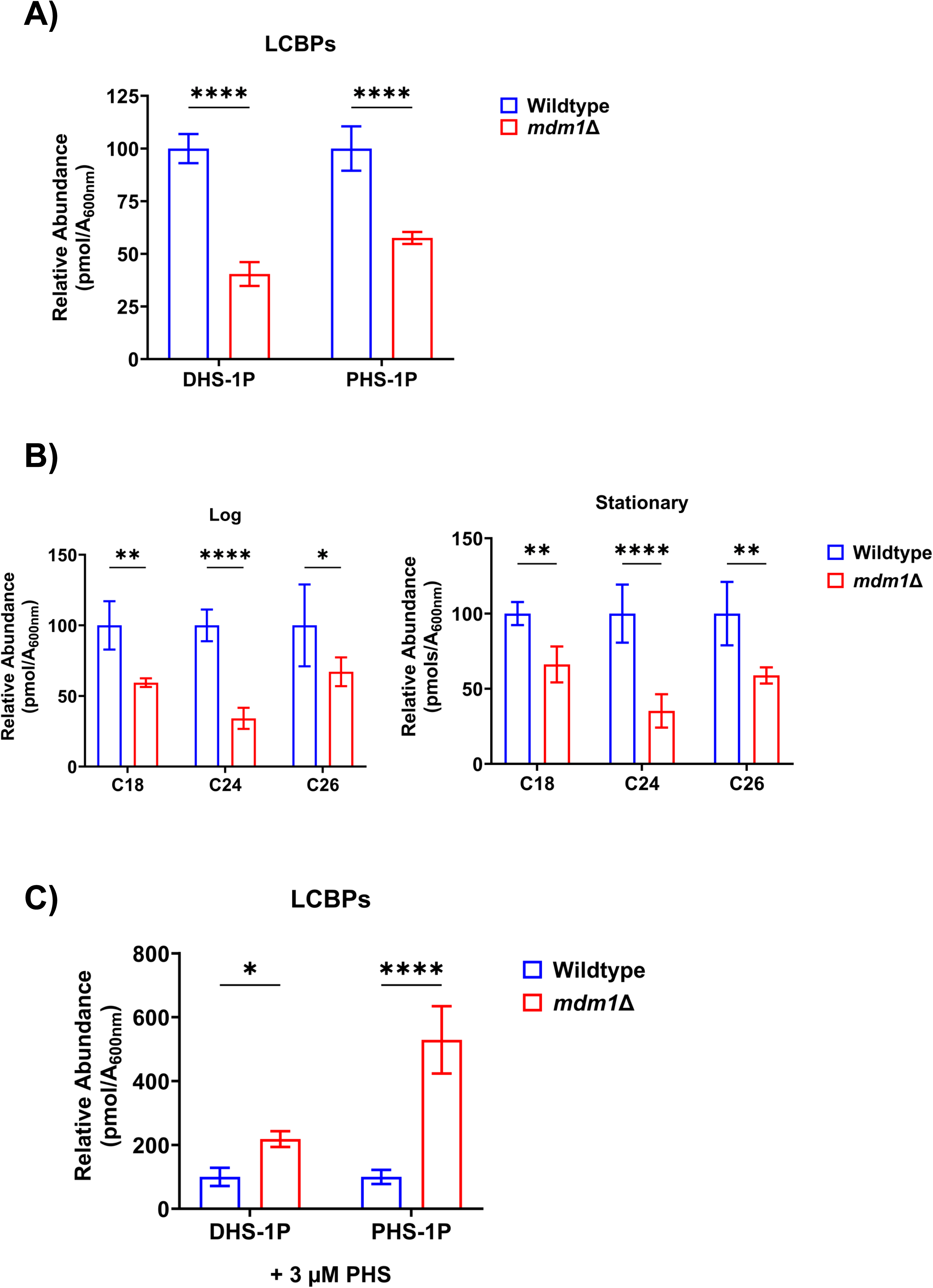
Loss of Mdm1 alters sphingolipid levels. A. Relative abundance of long-chain base phosphates (DHS-1P and PHS-1P) in wildtype and *mdm1*Δ cells grown to log phase in SCD media. Bars represent mean ± SD, n = 4, \*\*\*\**p* < 0.0001, Two-way ANOVA. B. Relative abundance of total sphingolipid by chain length (C18, C24, C26) in wildtype and *mdm1*Δ cells grown to log phase (left) or stationary phase (right) in SCD media (mean ± SD, n = 4, \**p* < 0.05, \*\**p* < 0.01, \*\*\*\**p* < 0.0001, two-way ANOVA). C. Relative abundance of long-chain base phosphates (DHS-1P and PHS-1P) in wildtype and *mdm1*Δ cells grown to log phase in media containing 3 μM PHS. Bars represent mean ± SD, n = 3, \*\*\*\**p* < 0.0001, Two-way ANOVA.

**Figure S2.**
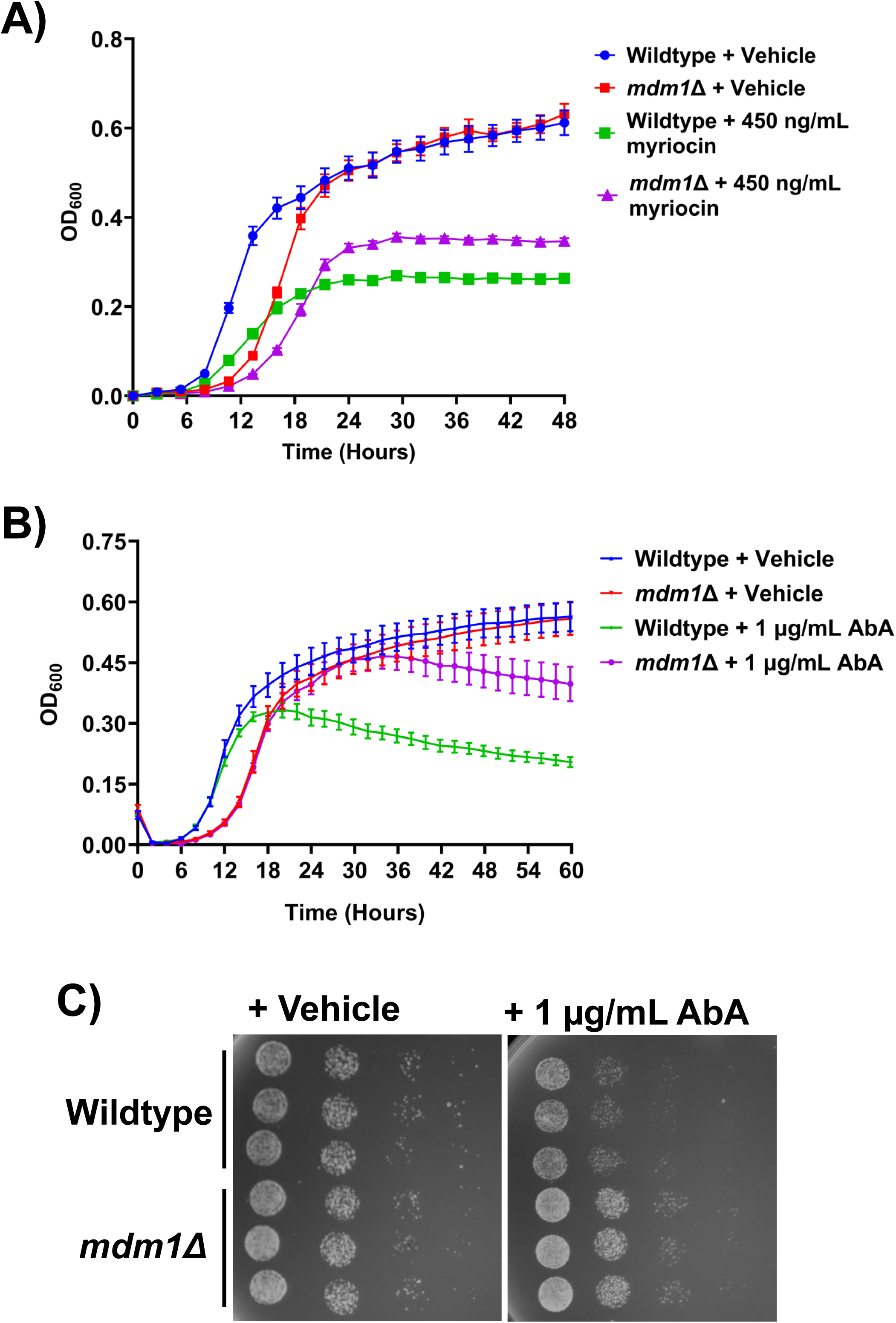
Loss of Mdm1 alters cellular adaptation to sphingolipid inhibitors. A. Growth curves of wildtype and *mdm1*Δ cells cultured in liquid medium with vehicle or 450 ng/mL myriocin. Optical density (OD₆₀₀) was monitored over time. Data represent mean ± SD, n = 4. B. Growth curves of wildtype and *mdm1*Δ cells cultured with vehicle or 1 µg/mL aureobasidin A (AbA). Optical density (OD₆₀₀) was measured over time. Data represent mean ± SD, n = 4. C. Serial dilution spotting assay of wildtype and *mdm1*Δ cells plated on medium containing vehicle or 1 µg/mL AbA. Cells were spotted in 5-fold serial dilutions and incubated at 30 °C. Images are representative of three independent experiments.

**Figure S3.**
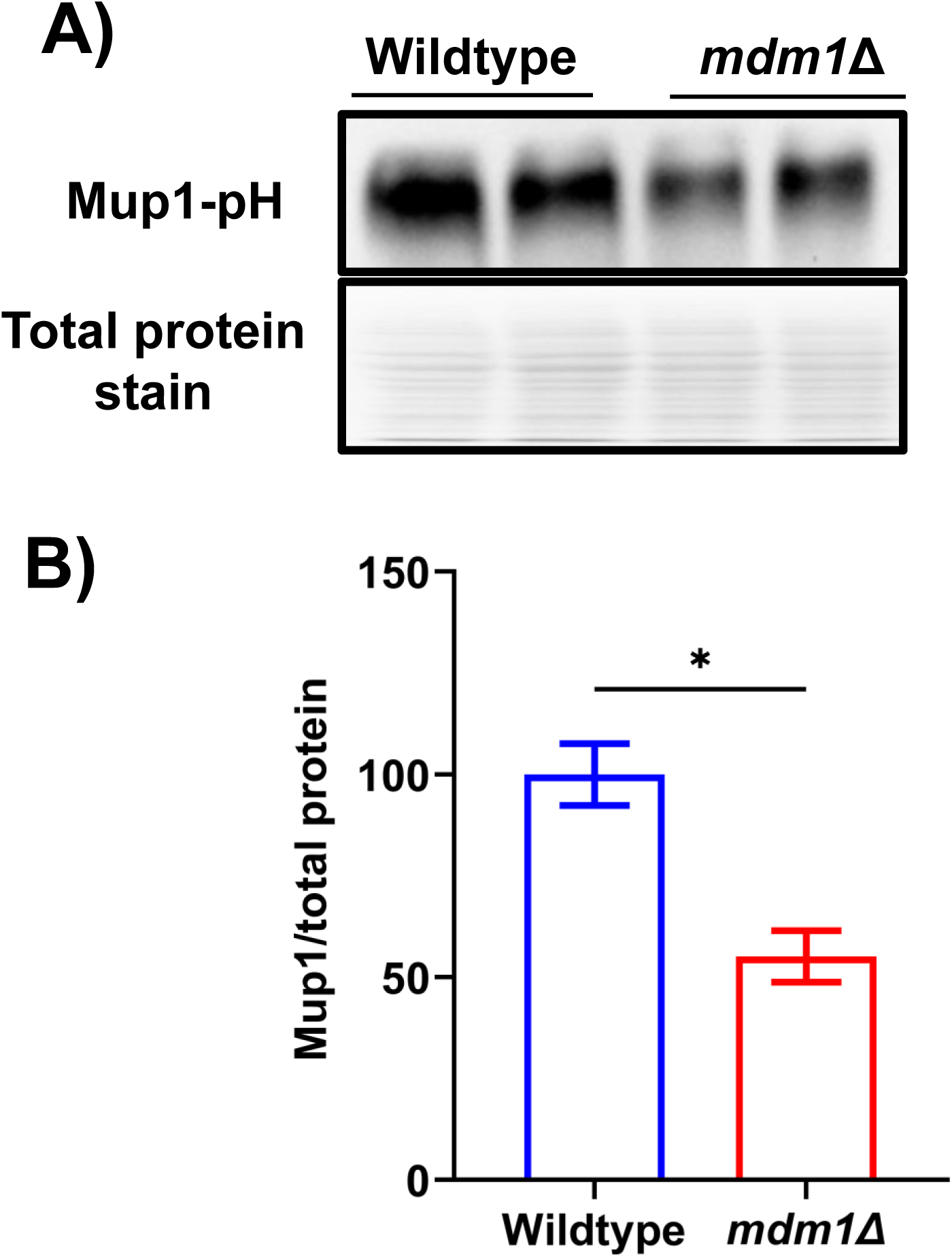
Reduced Mup1 abundance in *mdm1*Δ cells. A. Immunoblot analysis of Mup1-GFP levels in wildtype and *mdm1*Δ cells grown to log phase in −Met. Total protein stain is shown as loading control. B. Quantification of Mup1-GFP abundance from **A**, normalized to total protein. Bars represent mean ± SD, n = 2, \**p* < 0.05, students *t*-test.

**Figure S4.**
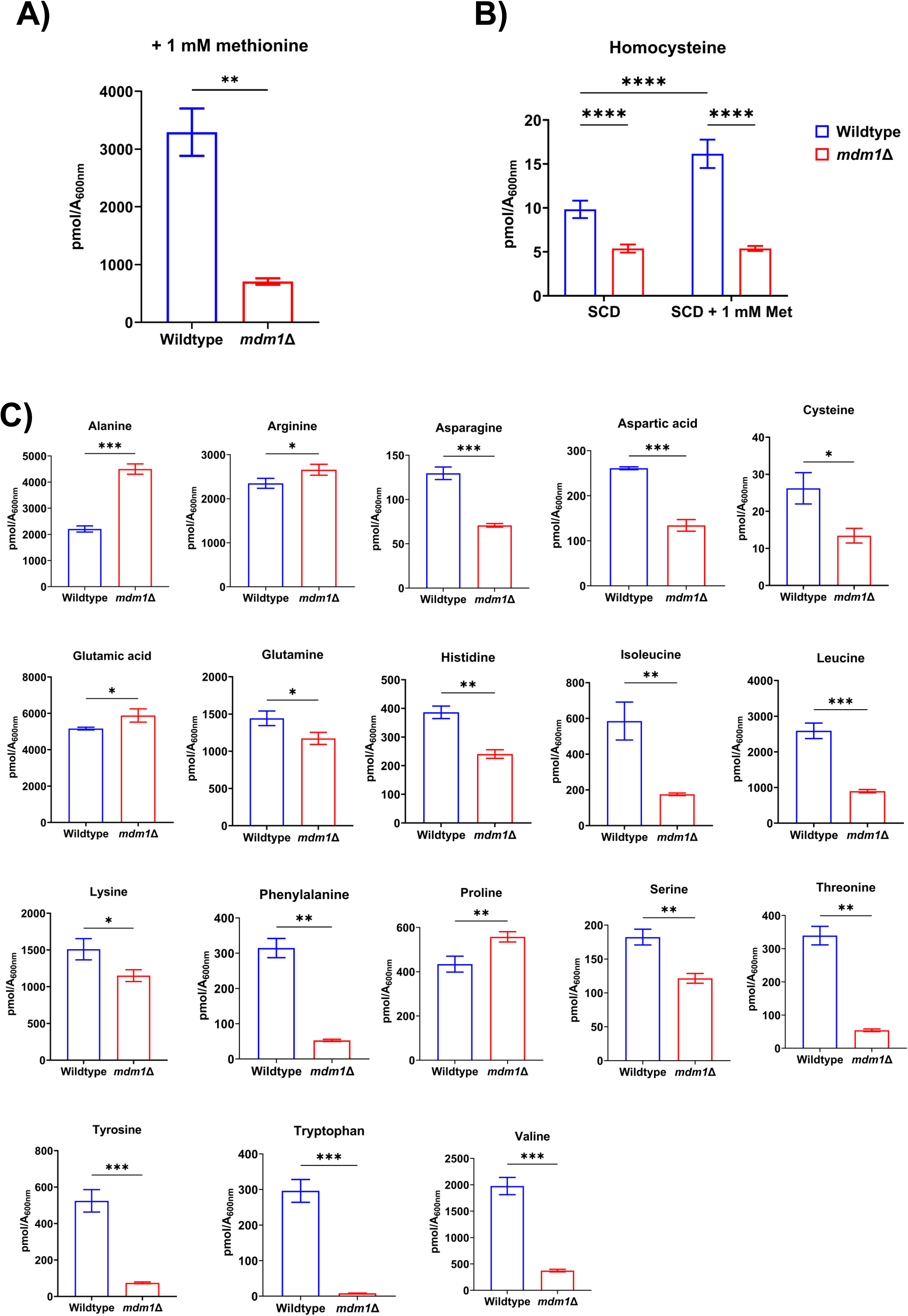
Methionine-cycle and amino acid profiling in *mdm1*Δ cells. A. Quantification of intracellular methionine levels in WT and *mdm1*Δ yeast supplemented with 1 mM methionine (mean ± SD, n = 3, \*\**p* < 0.01, students *t*-test). B. Intracellular homocysteine abundance in wildtype and *mdm1*Δ cells grown in SCD medium. Bars represent mean ± SD, n = 3, \*\**p* < 0.01, students *t*-test. C. Targeted metabolomic analysis of intracellular amino acids in wildtype and *mdm1*Δ cells grown in SCD. Data represent mean ± SD, n = 3; \**p* < 0.05, \*\**p* < 0.01, \*\*\**p* < 0.001 students *t*-test.

**Figure S5.**
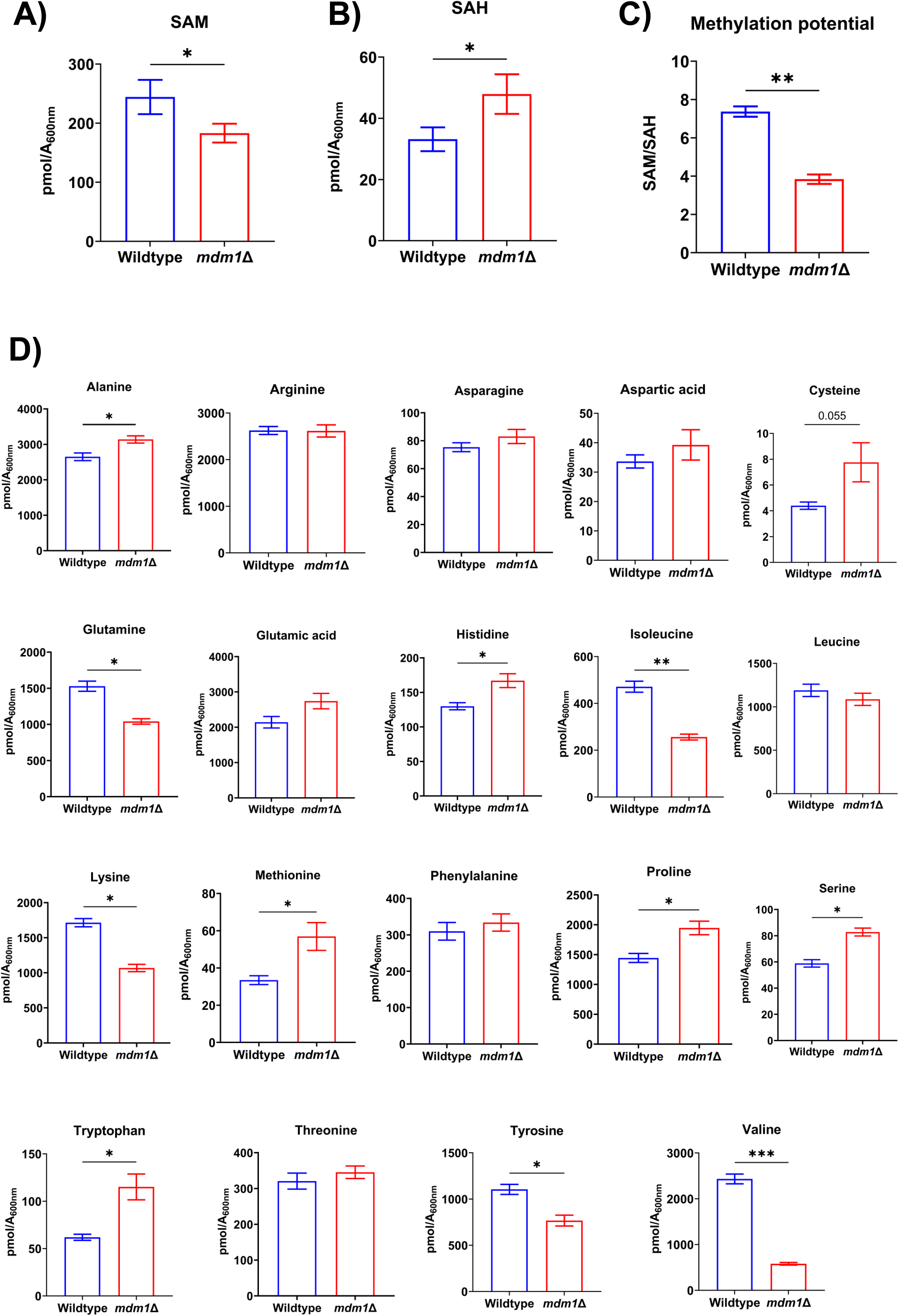
Effects of PHS treatment on methylation balance and amino acid pools. A. Intracellular SAM abundance in wildtype and *mdm1*Δ cells after PHS treatment. Bars represent mean ± SD, n = 3, \**p* < 0.05, students *t*-test. B. Intracellular SAH abundance in wildtype and *mdm1*Δ cells after PHS treatment. Bars represent mean ± SD, n = 3, \**p* < 0.05, students *t*-test. C. Methylation potential (SAM/SAH ratio) in wildtype and *mdm1*Δ cells following PHS supplementation (mean ± SD, n > 100 cells, \*\**p* < 0.01, students *t*-test). D. Targeted metabolomic analysis of intracellular amino acids in wildtype and *mdm1*Δ cells after PHS treatment. Data represent mean ± SD, n = 3; \**p* < 0.05, \*\*\**p* < 0.001 students *t*-test.

